# Human iPSC-Derived Cardiomyocytes Maladaptively Remodel to Cyclic Stretch in Hypoplastic Left Heart Syndrome

**DOI:** 10.64898/2026.10.11.758388

**Authors:** Riya S. Verma, Ivy Zhang, Xunyang Chen, Maddison S. Pierce, Ram Kumar Subramanyan, Megan L. McCain

## Abstract

**Background:** Hypoplastic Left Heart Syndrome (HLHS) is a severe congenital heart defect characterized by underdevelopment of left-sided cardiac structures, requiring staged surgical palliation that establishes the right ventricle as the single ventricle maintaining systemic circulation. After surgery, patients remain highly susceptible to heart failure, possibly due to the increased hemodynamic load imposed on the single surviving ventricle. Because the etiology of HLHS remains poorly understood, elucidating the mechanisms that drive ventricular dysfunction requires model systems that capture both patient-specific genetics and the dynamic mechanical environment of the ventricle. Thus, our objective was to compare how induced pluripotent stem cell (iPSC)-derived cardiomyocytes (iPSC-CMs) from patients with HLHS differ from control in static culture and in response to cyclic stretch.

**Methods:** Control iPSCs from two unaffected donors and iPSCs from four HLHS patients were differentiated into iPSC-CMs, engineered into aligned tissues, and cultured statically or exposed to cyclic 20% uniaxial stretch at 1 Hz for 48h. Sarcomere organization was quantified in static tissues using images of immunostained α-actinin. Calcium transients at 1 Hz pacing frequency in static and stretched tissues were evaluated using Fluo-4 calcium indicator dye. Bulk RNA sequencing was performed to evaluate stretch-induced transcriptomic changes in control and HLHS iPSC-CMs.

**Results:** In static culture, HLHS iPSC-CMs had modest defects in sarcomere organization and calcium transient morphology compared to control. In response to cyclic stretch, control iPSC-CMs exhibited calcium transients with quicker rise time, a response observed in only one of four lines of HLHS iPSC-CMs. Transcriptomic analysis revealed that all four lines of HLHS iPSC-CMs failed to upregulate calcium handling and sarcomere-associated genes in response to stretch, unlike control iPSC-CMs. Stretch also induced HLHS iPSC-CMs to exhibit pronounced activation of inflammatory signaling pathways, a response not observed in control iPSC-CMs.

**Conclusions:** iPSC-CMs from HLHS patients demonstrate aberrant responses to cyclic stretch, including blunted adaptation of calcium transients and heightened inflammatory signaling. These findings implicate defective mechanoadaptation as a potential contributor to single ventricle failure and identify potential candidate biomarkers and new opportunities for therapeutic interventions for HLHS patients.

## Introduction

Hypoplastic left heart syndrome (HLHS) is a single ventricle defect defined by an underdeveloped, non-functional left ventricle. HLHS has a prevalence of approximately 2.5 per 10,000 live births in the United States^1^ and is fatal without treatment. Patients with HLHS undergo three staged surgical procedures (Norwood, superior cavopulmonary connection, total cavopulmonary connection) that progressively reconfigure the circulation such that systemic venous return is passively directed to the pulmonary circuit, establishing the right ventricle as the pump for the systemic circulation^2,3^. Advances in pediatric cardiac surgery have substantially improved early survival in HLHS patients. However, patients have an ongoing risk for multiple morbidities, including ventricular dysfunction, arrhythmia, heart failure, and plastic bronchitis, leading to transplant free-survival rates at 35 years of age of approximately 30%^4^.

Advancing therapeutic strategies for HLHS is especially challenging because its etiology is extraordinarily complex and incompletely understood. Although several genes, such as *Notch1*, have been implicated in the development of HLHS-like morphology in animal models^5^, no single gene or combination of genes has been strongly correlated with the pathogenesis of HLHS in human patients. Thus, the ability to differentiate cardiac cells from induced pluripotent stem cells (iPSCs) derived from HLHS patients has been transformative for identifying abnormal phenotypes and disease mechanisms in a patient-relevant context^6^. Previous studies have demonstrated that iPSC-derived cardiomyocytes (iPSC-CMs) from HLHS patients exhibit dysregulated NOTCH signaling, impaired contractility, and mitochondrial energetics, suggesting that HLHS cardiomyocytes have cell-intrinsic defects^7–10^. However, a critical limitation of these studies is that cardiomyocytes were cultured as static, un-structured monolayers, a configuration that poorly mimics the dynamic biophysical microenvironment that these cells experience and must adapt to throughout all stages of life.

During embryonic development, hemodynamic loading and the resulting mechanical stretch of the primitive myocardium promotes the proliferation and maturation of cardiomyocytes, as demonstrated in chick and zebrafish^11–13^. In the mature heart, the amount of contractile force generated by the ventricle increases with the mechanical stretch it experiences during diastolic filling, as defined by the classical Frank-Starling Law. However, excessive stretch is often pathological, as in the case of volume overload, which stretches the ventricle to an abnormally high extent and induces chamber dilation that often progresses to heart failure^13^. These effects have also been replicated in human iPSC-CMs *in vitro*, which activate mature gene expression programs and exhibit enhanced sarcomere organization, calcium kinetics, and force generation in response to 5-10% cyclic stretch^14–18^. However, application of ≥25% cyclic stretch induces maladaptive remodeling by disrupting calcium handling and promoting structural disarray^16^. Thus, cardiomyocytes exquisitely sense and adapt to mechanical stretch in many contexts, which is key to the normal development and physiology of the heart. Notably, the surgical procedures for HLHS patients expose the right ventricle to profound alterations in hemodynamic loading, including periods of both volume and pressure overload followed by unloading^3^. Thus, defining how HLHS cardiomyocytes respond to stretch is especially important for establishing the mechanisms by which the single ventricle adapts or fails under increased hemodynamic demands, such as after surgical intervention. However, processes of mechanotransduction in HLHS CMs have not been thoroughly investigated.

We hypothesized that HLHS cardiomyocytes have impaired responses to mechanical loading that contribute to cardiac dysfunction. To test our hypothesis, we engineered aligned tissues from iPSC-CMs derived from four HLHS patients and two control donors and subjected them to 20% cyclic stretch for 48h. We then assessed calcium transients and transcriptional responses and compared them to tissues maintained in static culture. Control iPSC-CMs adopted a more mature calcium transient morphology in response to stretch, a response that was muted in HLHS iPSC-CMs. Cyclic stretch also induced control iPSC-CMs to up-regulate several key genes and pathways related to excitation-contraction coupling while HLHS iPSC-CMs instead up-regulated inflammatory and stress responses. These findings suggest that HLHS iPSC-CMs have a maladaptive response to mechanical loading, which may contribute to progressive ventricular dysfunction and abnormal heart development in HLHS patients.

## Materials and Methods

### Data Availability

The RNA sequencing data generated in this study are available in the Gene Expression Omnibus (GEO) under the identifier GSE347668.

### Cell Lines

iPSCs reprogrammed from two unaffected donors, Control 1 (ND03719; male, 33 years) and Control 2 (ND05280; female, 72 years), were generously provided by the lab of Justin Ichida at USC^19^. Four lines of HLHS iPSCs were obtained from the HLHS consortium. At birth, patient umbilical cord blood was obtained and isolated PBMCs were used to reprogram cells into iPSCs. Two male and two female lines of iPSCs from HLHS patients were used in this study. HLHS 1 (male) and HLHS 2 (female) both presented with sporadic classical HLHS with no family history of cardiac abnormalities. HLHS 3 (male) presented with classical HLHS and a paternal history of bicuspid aortic valve and dilated aorta, alongside a chromosome 2 duplication spanning 16 genes. HLHS 4 (female) presented with sporadic classical HLHS and a smaller chromosome 2 duplication spanning 3 genes.

### Maintenance of iPSCs and Differentiation to Ventricular CMs

iPSCs were thawed and seeded onto plates coated with Corning® Matrigel® hESC-Qualified Matrix (Corning, cat no. 354277; 1:100 in Gibco Knockout DMEM, ThermoFisher, cat no. 10829018; 1 hour at 37°C). Cells were initially cultured in mTeSR+ (STEMCELL Technologies, cat no. 100-0276) supplemented with 10 µM ROCK inhibitor Y-27632 (SelleckChem, cat no. S6390) for 24h, after which the media was replaced with mTeSR+ without Y-27632. Once cells reached 60-70% confluence, they were passaged by rinsing with PBS without Ca^2+^/Mg^2+^, followed by incubation in 0.5 µM UltraPure™ EDTA (ThermoFisher, cat no. 15575020) for 5 minutes at 37°C. EDTA was then aspirated, and cells were detached using a serological pipette. The cell suspension was split at a 1:6-1:3 ratio onto Matrigel-coated plates. All iPSCs were kept under P30 and all lines were routinely screened for mycoplasma using Universal Mycoplasma PCR Detection Kit (ATCC, cat no.30-1012K) prior to seeding for differentiations.

For differentiation to CMs (Supplemental Figure 1A), iPSCs were singularized by incubating with Accutase^TM^ (STEMCELL Technologies, cat no. 07920) for 5 minutes at 37°C. A P1000 pipette was used to triturate the cells and then neutralized with 1.5x volume of mTESR+ media. Cells were centrifuged at 300g for 5 minutes and the pellet was resuspended in mTESR+ media with 10µM Rock inhibitor Y-27632 and CloneR™2 (STEMCELL Technologies, cat no. 100-0691). Cells were counted and seeded at 0.75-2 x 10^6^ cells per well of a 6-well plate for differentiation. Ventricular CMs were then derived using STEMdiff™ Ventricular CM Differentiation Kit from (STEMCELL Technologies, cat no. 05010) according to manufacturer’s instructions. Cells typically started beating between day 8-10.

After day 10-12, cardiomyocytes were purified and matured using glucose deprivation and fatty acid supplementation (Supplemental Figure 1B), similar to previous protocols^20^. Briefly, cells were cultured in RPMI without Glucose (ThermoFisher, cat no. 11879020) with B27 minus vitamin A (ThermoFisher, cat no. 12587010) for 48 hours. Cells were dissociated with 0.25% Trypsin-EDTA (ThermoFisher, cat. no. 25200056) for 10 minutes at 37°C, triturated using a P1000 pipette, neutralized with STEMdiff™ CM Support Medium (STEMCELL Technologies, cat. no. 05027) and passaged at a 1:3 ratio onto Matrigel-coated plates. 100 µL aliquots of cell suspension were removed for flow cytometry profiling, as described below. After 24h, media was replaced with RPMI without Glucose, containing B27 without vitamin A and fatty acids modified from a previous protocol^20^. The fatty acid media contained 25 µM Bovine Serum Albumin (BSA)-Palmitate Saturated Fatty Acid Complex (Caymen Chemical, cat no.2 9558), 10 µM of BSA-Linoleic and BSA-Oleic Acids (Millipore Sigma, cat no. L9530-5ML, cat no. O3008-5ML), and 120 µM of L-carnitine (ThermoFisher, cat no. A17618-09). After 48h, the media was exchanged to 75% fatty acid media and 25% CM Maintenance media (STEMCELL Technologies) to balance fatty acid β-oxidation and glucose oxidation^21^. Cells were cryopreserved at day 17-19 using the STEMdiff™ CM Freezing Medium (STEMCELL Technologies, cat no. 05030) according to the manufacturer’s protocol. As needed, CMs were thawed in 6-well plates in STEMdiff™ CM Support Medium and kept in 75% Fatty Acid media for 48h for recovery prior to experiments. iPSC-CMs at day 20+ were used for all experiments (Supplemental Figure 1C).

### Flow Cytometry

For each differentiation, ∼1x10^5^ dissociated cells were used to evaluate the differentiation efficiency based on quantification of cardiac troponin T (cTnT) expression by flow cytometry. Cells were pelleted and fixed for 10 minutes at room temperature with 1mL of 4% Pierce™ Formaldehyde (w/v), Methanol-free (ThermoFisher, cat no. 28908), diluted in PBS Ca^-^/Mg^-^. Fixed cells were washed three times with PBS Ca-/Mg- and centrifuged at 300g. Cell permeabilization was performed in FACS buffer containing 0.5% (w/v) saponin (ThermoFisher, cat no. A18820.22) and 4% FBS in PBS for 15 minutes at room temperature. Cells were incubated with cTnT monoclonal antibody (1:100; ThermoFisher, cat. no. MA5-12960) or Mouse IgG1 kappa isotype control (1:100) for 1h at room temperature, followed by three washes with PBS. Samples were next incubated for 1h at room temperature with goat anti-Mouse IgG (H+L) Cross-Adsorbed Secondary Antibody, Alexa Fluor™ 568 (ThermoFisher, cat no. A-11004) (1:100) and 1 µg/mL DAPI (4’,6-Diamidino-2-Phenylindole, Dilactate) (ThermoFisher, cat no. D3571). Samples were then washed three times and analyzed on the Beckmann Coulter CytoFLEX machine. Gates for cTnT+ were drawn based on negative IgG control (Supplemental Figure 1D).

### Fabrication of Micropatterned Coverslips and iPSC-CM Seeding

Cleanroom photolithography was used to make wafer templates for microcontact printing, as previously described^22,23^. Briefly, 4” silicon wafers were spin-coated with SU-8 2005 (Kayaku, cat no. NC9463827) or 3005 (Kayaku, cat no. Y311049) photoresist, soft baked, and exposed to UV light through a chrome photomask with 30 µm-wide lanes separated by 30 µm spacing using a mask aligner (MA BA6 Gen4). Wafers were then hard baked, soaked in developer solution to remove uncured photoresist, rinsed with 100% isopropanol, and dried. Wafers were silanized overnight in a desiccator with 3-Aminopropyltriethoxysilane (APTES). Polydimethylsiloxane (PDMS) (Dow Corning Sylgard 184, cat no. 4019862) was prepared and poured onto wafers, desiccated, and placed in a 65°C oven. The cured PDMS was then carefully removed from the wafer and manually cut into stamps for microcontact printing.

Stamps were sonicated in 70% ethanol for 30 minutes, then transferred into a biosafety cabinet, dried with compressed air, and coated with 50 µg/mL fibronectin (Corning, cat no. 47743-654) for 1 hour at room temperature. 18 mm glass coverslips spin-coated with PDMS were treated with ultraviolet-ozone (UVO) for 8 minutes (Jelight, model 342). Excess fibronectin was removed from stamps using compressed air and stamps were gently inverted onto UVO-treated PDMS-coated coverslips to transfer the fibronectin pattern. Coverslips were immediately placed in a 12-well plate, washed three times with PBS, and stored at 4°C with Gibco™Antibiotic-Antimycotic (100X) (ThermoFisher, cat no. 15240062). Before cell seeding, plates were placed without lids in the biosafety cabinet and treated with the UV sterilization bulb for 20 minutes.

Day 20+ iPSC-CMs were dissociated using STEMdiff™ Cardiomyocyte Dissociation Kit (STEMCELL Tech, cat no.05025) according to manufacturer’s instructions. Cells were counted using trypan blue and seeded at 1–1.4 x 10^5^ cells/cm^2^ onto micropatterned coverslips in STEMdiff™ Cardiomyocyte Support Medium (StemCell Tech, cat no. 05027). After 24 hours, media was exchanged to 75% fatty acid media. After an additional 24h, cells were used for downstream experiments.

### Fabrication of Micropatterned Stretchable Substrates and iPSC-CM Seeding

To micropattern stretchable membranes (Supplemental Figure 2A), gloss silicone sheeting of 0.01" thickness (Specialty Manufacturing) was laser-cut into 35 mm x 20 mm rectangles, sonicated in 90% ethanol for 30 minutes, and dried at 65°C for 20 minutes on a Petri dish covered in masking tape. Dried substrates were treated with UVO for 8 minutes and microcontact printed with fibronectin, using the protocol described above. In parallel, the bottom surfaces of 12 mm x 12 mm stretchable culture chambers (CuriBio) were plasma treated for 3 minutes and inverted onto the micropatterned silicone membrane. A small weight was placed on top of the chamber for 20 minutes to facilitate bonding. The bonded chambers were rinsed 3X with PBS without Ca^2+^/Mg^2+^, filled with Gibco™Antibiotic-Antimycotic (100X) (ThermoFisher, cat no.15240062), and sterilized under the UV light in the biosafety cabinet for 20 minutes. Prior to cell seeding, chambers were treated with 1% w/v Pluronic F-127 in deionized water for 5 minutes.

Day 20+ iPSC-CMs were dissociated using STEMdiff™ Cardiomyocyte Dissociation Kit (STEMCELL Tech, cat no.05025) according to manufacturer’s instructions and seeded onto micropatterned stretchable membranes at 3.5x10^5^ cells/cm^2^ in STEMdiff™ Cardiomyocyte Support Medium (StemCell Tech, cat no. 05027). After 24h, media was exchanged to 75% fatty acid media. Stretchable constructs were then loaded onto the CuriBio Cytostretcher and subjected to 20% strain at 1 Hz frequency (0.5 seconds of stretch followed by 0.5 seconds relaxation) for 48h, followed by downstream experiments. In parallel, tissues on stretchable membranes were maintained in static culture for the same duration.

### Sarcomere Immunohistochemistry and Analysis

Coverslips were fixed in 4% paraformaldehyde (ThermoFisher, cat no. 28908) for ten minutes with gentle rocking, then rinsed 3X with ice cold PBS with Ca^2+^/Mg^2+^ for five minutes. Cells were next permeabilized in 0.1% Triton X-100 in Hanks Balanced Salt Solution (HBSS) for 10 minutes with gentle rocking, followed by 3X 5-minute rinses in HBSS. Coverslips were then incubated in a blocking solution of 3% bovine serum albumin in PBS for 1h on a rocker at room temperature, followed by primary antibodies (mouse anti-cardiac troponin T, ThermoFisher, MA5-12960; rabbit anti-α-actinin, Thermofisher, 701914; rabbit anti-fibronectin, Sigma Aldrich, F3648) diluted 1:200 in 3% BSA in PBS and incubated at 4°C on a rocker overnight. After 12h, cells were rinsed 3X with PBS for 5 minutes and incubated with 1:1000 Alexa Fluor™ cross-adsorbed secondary antibodies (Goat anti-Rabbit IgG (H+L) Cross-Adsorbed Secondary Antibody, Alexa Fluor™ 488, ThermoFisher, A-11008; Goat anti-Mouse IgG (H+L) Cross-Adsorbed Secondary Antibody, Alexa Fluor™ 568, ThermoFisher, A-11004) in PBS in the dark for one hour with rocking at room temperature. Coverslips were rinsed 3X with PBS for 10 minutes, after which coverslips were inverted onto 7 µL of DAPI Fluoromount-G (Southern Biotech, cat. no. 0100-20) and mounted onto glass slides. Images of α-actinin were acquired using Zeiss LSM 880 Confocal Laser Scanning Microscope with a 63x water and at least five regions with two lanes of cardiomyocytes each were imaged per coverslip.

Sarcomere length and organization were quantified using the SotaTool software package^24^. Briefly, each image file was opened in ImageJ and cropped to isolate individual lanes (two per file). Z-stack projections were generated and exported for analysis. SotaTool uses a gray-level co-occurrence matrix (GLCM)-based texture analysis that computes pixel-intensity correlation across 180 angles over a range of pixel offsets^24^. Striation patterns, such as sarcomeres, produce a damped oscillating correlation curve. The algorithm automatically selects the angle of maximum correlation to determine sarcomere length. Sarcomere organization is quantified as the height of this maximum peak above its preceding minimum. An alignment index is derived from the coefficient of variation across all angular curves at that offset. For each coverslip, the values across at least five regions were averaged as a single data point. For each patient line, we collected data from at least two independent differentiations.

### Live Calcium Imaging and Analysis

iPSC-CMs on coverslips or stretchable membranes were incubated in 0.5 mg/mL Fluo-4, AM (ThermoFisher, cat no. F14201) in Pluronic F-127 20% solution in DMSO (ThermoFisher, cat no. P3000MP) resuspended in 1 mL of Dulbecco’s Modified Eagle Medium (DMEM) for 15 minutes at 37°C. Coverslips or membranes were then rinsed in DMEM and transferred to the stage of a Nikon Ti inverted fluorescence microscope with an incubator set to 32°C. A platinum field electrode was inserted into the media to pace tissues at 1 Hz with a pulse duration of 14 ms (Supplemental Figure 2B). Images were acquired with a 10x objective and an Andor Zyla scientific CMOS camera at 100 frames per second (10 ms exposure) for a 3s interval. One region was imaged per sample. Each region contained 50–75 micropatterned lanes of iPSC-CMs.

For calcium imaging analysis (Supplemental Figure 3A), the brightest frame (maximum summed intensity) was identified and intensity for the video was rescaled to the 2nd–98th percentile range. A Butterworth filter, Gaussian smoothing (σ = 1.5), and Otsu thresholding was used to generate a binary mask, followed by removal of small objects (<5000 pixels) to distinguish regions with and without iPSC-CMs and isolate individual lanes of cells. Individual lanes were then identified as connected components using a breadth-first search over 8-connected neighborhoods. The mean pixel intensity within each lane was extracted across all frames to yield 50-75 lanes of calcium traces indexed to time (100 fps). A csv containing transients of lanes was then analyzed using a modified version of previous software^25^. Per-lane calcium traces were normalized (F/F_0_) and analyzed to identify candidate peaks (local maxima) (Supplemental Figure 3B). Retained peaks were further refined to eliminate spurious or duplicate maxima within each transient. For each transient, amplitude, peak timing, rise and decay kinetics (time to 50/85/90% of peak), downstroke velocity, beat period, and beat frequency were quantified, along with single- and double-exponential decay time constants fit to the relaxation phase. The full code for image lane isolation can be found here: https://github.com/RiyaS-Verma/CalciumLaneAnalysis. Calcium dynamics were then analyzed for each lane using a modified script that can be accessed here: https://github.com/RiyaS-Verma/multilineage-organoid_RV. Each data point is a coverslip or membrane containing an average of 57 lanes of iPSC-CMs. For each patient line, we collected data from at least two independent differentiations.

### RNA Sequencing and Analysis

For each iPSC line, two independent differentiations of iPSC-CMs were pooled and seeded at 3.5 x 10^5^ cells/cm^2^ onto six micropatterned stretchable membranes. For each line, three chambers were cultured statically and three were stretched for 48h, as described above. Tissues were then lysed immediately using the lysis buffer from Qiagen’s RNeasy Mini Kit (Qiagen, cat no. 74104) supplemented with 10 µL β-mercaptoethanol (β-ME). Cells were incubated in lysis buffer for 5 minutes and triturated to fully dissociate them from the membrane. Lysates were homogenized by passage through a 21-gauge needle. Total RNA was isolated using the RNeasy Mini Kit according to the manufacturer’s instructions. All 36 samples successfully yielded RNA (10.7–26.7 ng/µL). Samples underwent RNA purification and paired-end sequencing at Novogene on the Illumina NovaSeq X Plus Series (PE150). See supplemental materials for detailed RNA analysis.

### Statistical Analysis

Unless otherwise noted, data are presented as mean ± standard deviation (SD) with a minimum sample size of n=5. All conditions contained at least two independent differentiations per sample. One-way ANOVA with a Tukey’s multiple comparison was used to analyze datasets that included more than two experimental groups. Comparisons between two variables were performed using an unpaired t-test with a Welch’s correction, as SD could not be assumed to be equal. Cohen’s d was calculated as d = (μ₂ − μ₁)/σ, where µ are the means of the respective groups and σ is the pooled standard deviation. Cohen’s U₃ was calculated as U₃ = Φ(d) and the overlapping coefficient as OVL = 2Φ(−d/2), where Φ is the cumulative distribution function of the normal distribution. Stretched and non-stretched tissues were compared as independent groups; n is defined as each discrete tissue and p<0.05 was considered significant. Analyses were performed in Python using NumPy, SciPy, and Matplotlib.

## Results

### Micropatterned iPSC-CMs Reveal Altered Sarcomere Organization and Relatively Normal Calcium Dynamics in HLHS

iPSCs from two male and two female HLHS patients, including one male and one female with chromosome 2 duplications (dup(2)), were differentiated into ventricular CMs. We refer to the male and female HLHS non-dup(2) iPSC-CMs as HLHS Line 1(M) and 2(F), respectively, and the male and female HLHS dup(2) iPSC-CMs as HLHS Line 3(M) and 4(F), respectively. We also differentiated iPSCs from male and female control donors, referred to as Control 1(M) and 2(F), respectively. Control and HLHS lines exhibited spontaneous rhythmic contractions and similar differentiation efficiencies, as quantified using flow cytometry for cardiac troponin (cTnT) (**Figure 1A**). To assess differences in sarcomere structure, we seeded iPSC-CMs onto 30 µm-wide micropatterned lanes of fibronectin (**Figure 1B**) and immunostained tissues for sarcomeric α-actinin (**Figure 1C**). Sarcomeres were significantly longer in Control 1(M) (1.6 ± 0.07 µm) compared to HLHS 1(M) (1.4 ± 0.06 µm; *p=0.0006*) and HLHS 2(F) (1.5 ± 0.02 µm; *p=0.023*) (**Figure 1D**), suggesting modest deficits in sarcomere structure in two HLHS lines. Sarcomere organization showed a more consistent and broader pattern of disruption across HLHS lines (**Figure 1E**). Control 1(M) had the highest organization index (0.1 ± 0.01), which was significantly greater than all four HLHS lines (HLHS 1(M), *p=0.0007*; HLHS 2(F), *p=0.0048*; HLHS 3(M), *p=0.018*; HLHS 4(F), *p=0.018*). Control 2(F) (0.1 ± 0.02) also had significantly greater organization index compared to HLHS 1(M) (*p=0.033*). All four HLHS lines had statistically similar sarcomere length and organization compared to each other. Together, these findings demonstrate that HLHS iPSC-CMs exhibit some impairment in sarcomere assembly compared to control iPSC-CMs.

**Figure 1:**
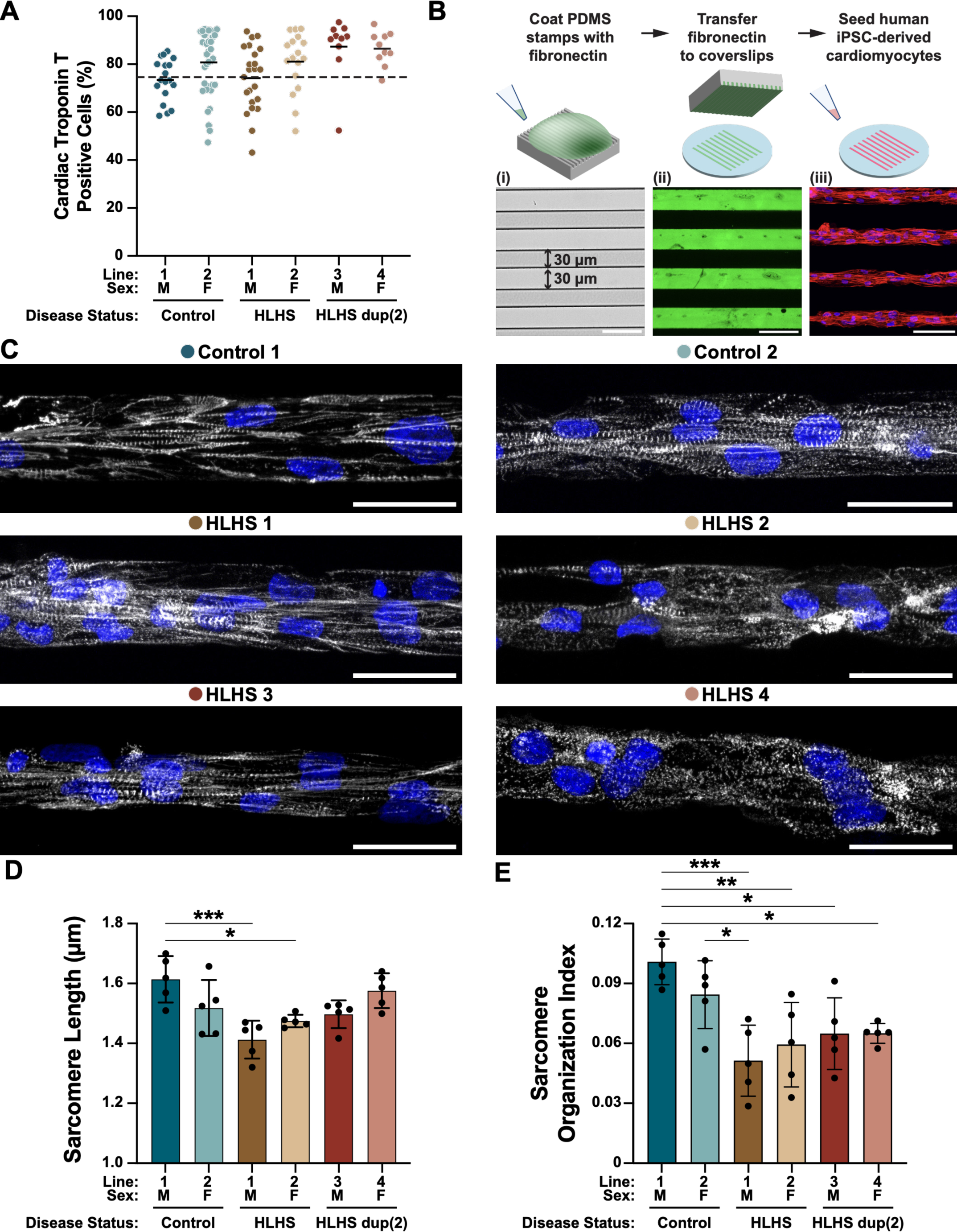
Sarcomere structure in control and HLHS iPSC-CMs. (A) Percentage of cardiac troponin T-positive (cTnT+) cells after differentiation to iPSC-CMs for each indicated line. Each point represents a sample from an independent differentiation batch. Dashed line indicates 75% CMs, the threshold used to select batches for downstream experiments. Solid bars represent mean values per line. Differentiation efficiency did not significantly differ between any lines based on one-way ANOVA and Tukey’s multiple comparisons test (*p*>0.05), n=10-36 differentiations per line. (B) Top: schematic of microcontact printing. Bottom: (i) brightfield image of PDMS stamp features, (ii) micropatterned fibronectin lanes (green: fibronectin), and (iii) iPSC-CMs cultured on micropatterned fibronectin lanes (red: cTnT, blue: DAPI). Scale bars = 60 µm. (C) Representative images of micropatterned iPSC-CMs from each indicated line (white: α-actinin, blue: DAPI). Scale bars = 30 µm. (D) Sarcomere length and (E) sarcomere organization index for iPSC-CMs from each indicated line. Each point represents the average of five regions collected from a single coverslip, collected from at least two independent differentiation cycles. Bars indicate mean±SD, n=5 per line. Statistical significance determined by one-way ANOVA with Tukey’s multiple-comparisons test (\**p*<0.05, \*\**p*<0.01, \*\*\**p*<0.001).

We next evaluated if HLHS iPSC-CMs exhibit defects in calcium handling as a functional metric. We incubated micropatterned 30-µm wide lanes of control and HLHS iPSC-CMs with Fluo-4 and imaged tissues using high-speed fluorescence microscopy (Supplemental Video S1 and S2). Spontaneous beat rate was highly variable within and across lines and did not differ significantly between control and HLHS lines (Supplemental Figure 3C). To compare calcium transients, we used field stimulation to electrically pace tissues at 1 Hz (**Figure 2A**). All control and HLHS iPSC-CMs successfully responded to 1 Hz pacing (Supplemental Figure 3D). The time to 50% peak calcium (**Figure 2B**) was significantly lower in Control 2(F) compared to all four HLHS lines (HLHS 1(M), *p=0.046*; HLHS 2(F), *p<0.0001*; HLHS 3(M), *p=0.0013*; HLHS 4(F), *p=0.011*). The time to 90% decay (**Figure 2C**) was significantly higher in Control 2(F) compared to three HLHS lines (602.2 ms vs. 471.3–475.4 ms; HLHS 1(M), *p=0.048*; HLHS 3(M), *p=0.042*; HLHS 4(F), *p=0.039*). These data suggest that calcium handling kinetics were generally faster in Control 2(F) compared to all HLHS lines. However, calcium dynamics were not significantly different between Control 1(M) and the four HLHS lines, suggesting that HLHS lines are still within relatively normal ranges. The inter-line variability, including between the two control lines, is consistent with the reported heterogeneity of iPSC-CMs.

**Figure 2:**
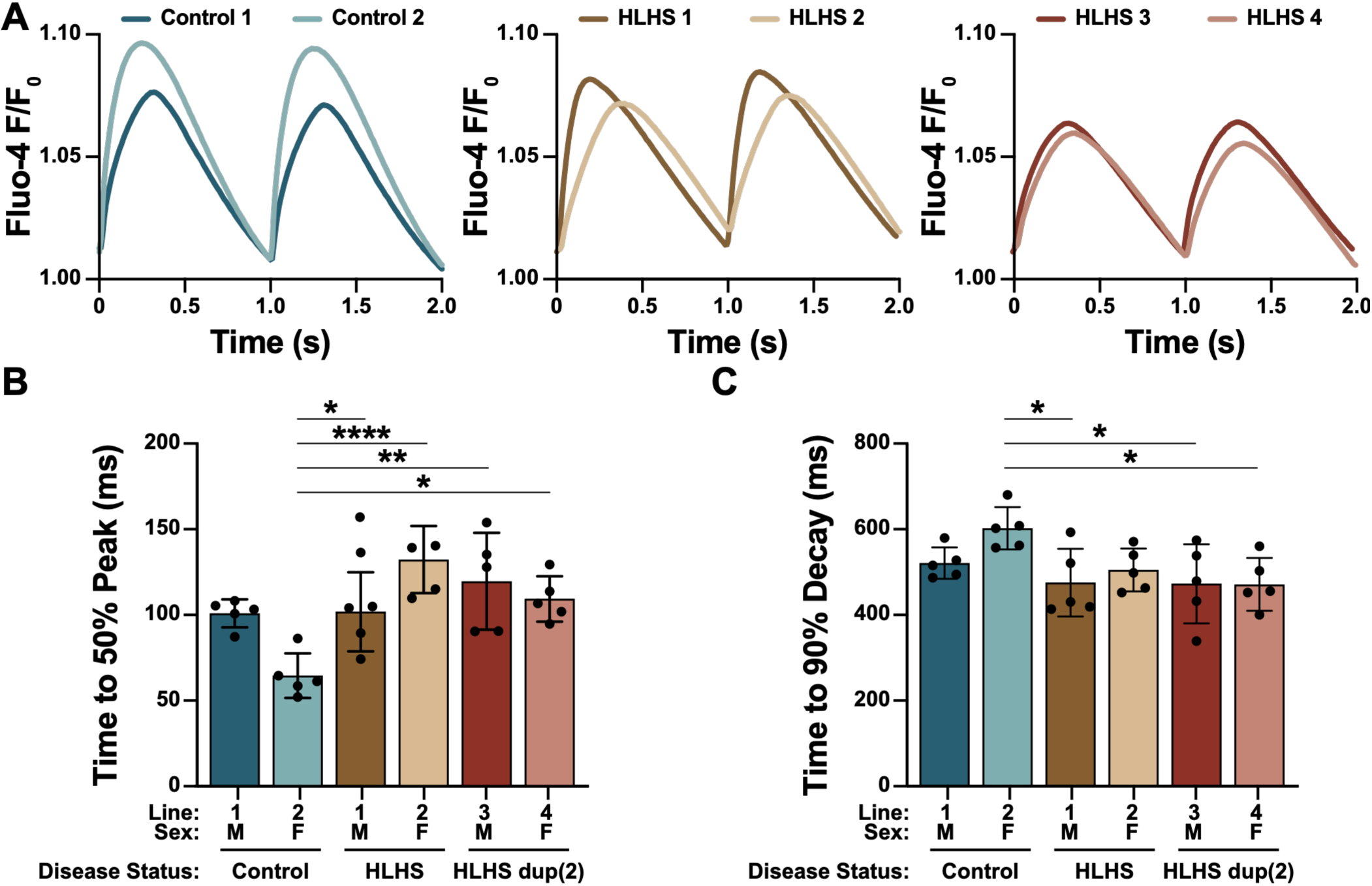
Calcium transients in control and HLHS iPSC-CMs. (A) Representative calcium transients at 1 Hz pacing for iPSC-CMs from each indicated line. Each trace is the average of 15 transients per line. (B) Time to 50% peak calcium and (C) time to 90% calcium decay for iPSC-CMs. Each point represents the average of five independent coverslips, containing 33-89 lanes, collected from at least two independent differentiation cycles. Bars indicate mean±SD, n=5 per line. Statistical significance determined by one-way ANOVA with Tukey’s multiple-comparisons test (\**p*<0.05, \*\**p*<0.01, \*\*\**p*<0.001, \*\*\*\**p*<0.0001).

### Calcium Transients in HLHS and Control iPSC-CMs Differentially Respond to Cyclic Stretch

Because adaptation of CMs to hemodynamic load is critical for normal cardiac development and physiology, we next asked if HLHS iPSC-CMs are defective in sensing or responding to mechanical stretch. To test this, we micropatterned 30-µm wide lanes of iPSC-CMs on stretchable silicone membranes and subjected them to 48h of 1 Hz cyclic uniaxial 20% stretch or static culture, followed by Fluo-4 imaging in electrically paced tissues (Supplemental Video S3-S6). Across all lines, the amplitude of peak Fluo-4 F/F_0_ trended lower after stretch and reached significance in Control 1(M) (**Figure 3A**, Supplemental Figure 4A), an effect that has also been observed in neonatal rat ventricular myocytes^26^. However, the change in amplitude is difficult to interpret because Fluo-4 is non-ratiometric.

**Figure 3:**
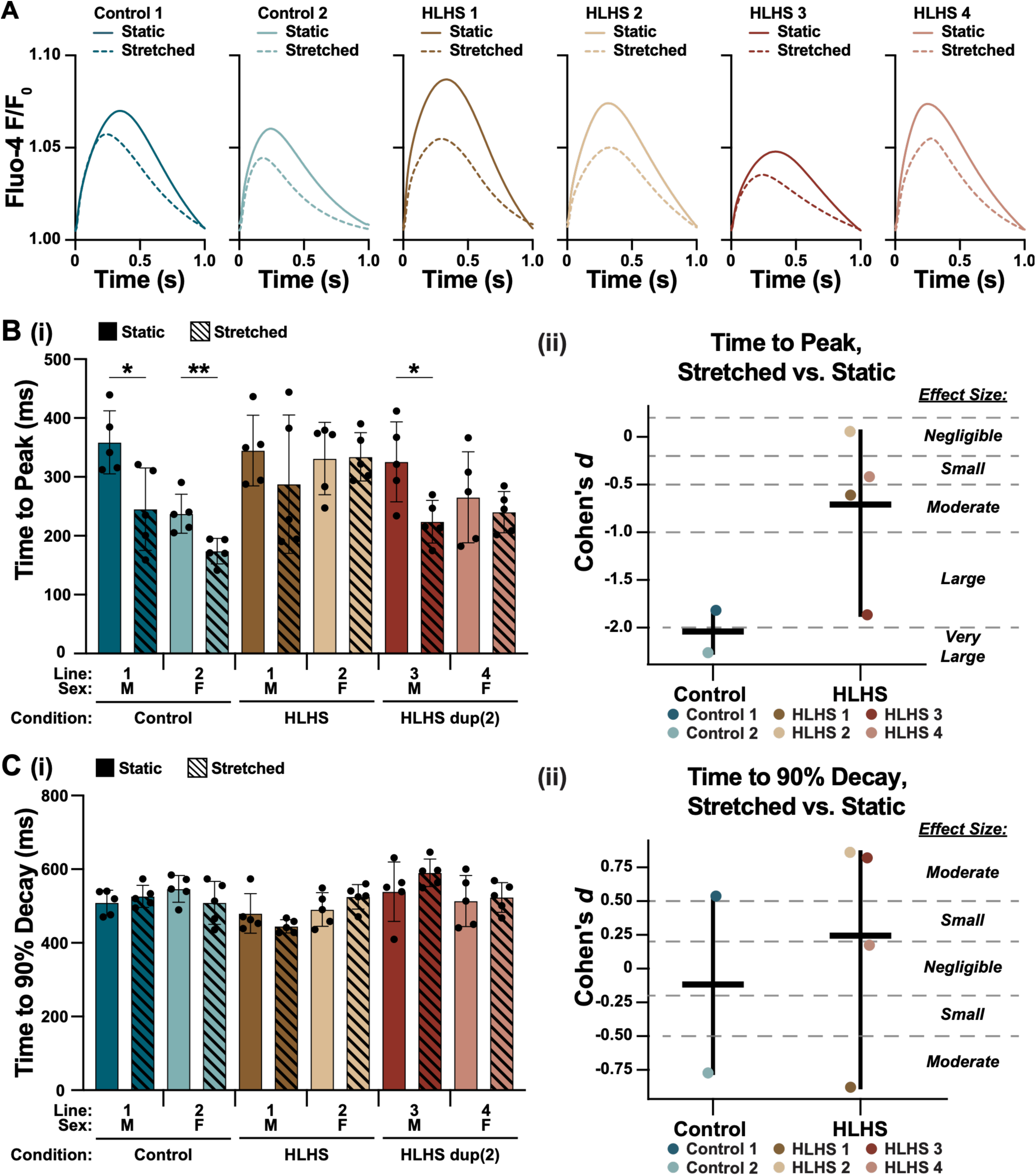
Stretch-induced remodeling of calcium transients in control and HLHS iPSC-CMs. (A) Representative calcium transients at 1 Hz pacing for iPSC-CMs from each indicated line. Solid lines indicate static tissues, and dashed lines indicate stretched tissues. Each trace is an average of 189-312 regions of iPSC-CMs per line, each collected from 5 tissues on stretchable membranes. (B) Time to peak calcium and (C) time to 90% calcium decay for iPSC-CMs for each indicated line and stretch condition. For (i), each point represents the average of multiple regions collected from a single tissue on a stretchable membrane, collected from at least two independent differentiation cycles. Bars indicate mean±SD, n=5 per line. Statistical significance between static and stretched tissues for each line determined by Welch’s t-test (\**p*<0.05, \*\**p*<0.01, \*\*\**p*<0.001, \*\*\*\**p*<0.0001). (ii) Cohen’s *d*, comparing stretched versus static with pooled standard deviation, for each indicated line, grouped by control or HLHS. Horizontal bars denote the mean and vertical bars denote the range. Dotted lines denote boundaries between relative effect sizes.

In terms of calcium dynamics, exposure to stretch significantly reduced the time to peak calcium in both Control 1(M) (358.7 vs. 245.3 ms; *p=0.022*) and Control 2(F) (237.4 vs. 174.1 ms; *p=0.0092*) (**Figure 3B**). In contrast, three of four HLHS lines showed no significant change in time to peak calcium due to stretch (HLHS 1 (344.8 vs. 287.8 ms; *p=0.372*), HLHS 2 (331.2 vs. 334.2 ms; *p=0.932*), and HLHS 4 (265.3 vs. 240.2 ms; *p=0.535*)). Only HLHS 3(M) exhibited a significant reduction in time to peak due to stretch (325.6 vs. 224.0 ms; *p=0.025*). To compare effect sizes of stretch, we calculated Cohen’s d for the stretch response of each line (**Figure 3B**). Control 1(M) and Control 2(F) demonstrated a large (d=1.84) and very large (d=2.23) effect size, respectively. HLHS 3(M) demonstrated a large effect size (d=1.87) but the three other HLHS lines demonstrated a negligible to moderate effect size (d = 0.06–1.87). Similar trends were observed in time to 50% peak calcium (Supplemental Figure 4B). Time to 90% decay (**Figure 3C**) and 50% decay (Supplemental Figure 4C) were largely unaffected in all control and HLHS lines, with no significant differences due to stretch and negligible to moderate effect sizes across all lines. These data indicate that control iPSC-CMs consistently adapt to stretch by accelerating calcium release but most lines of HLHS iPSC-CMs lack this response.

### Conserved Transcriptional Programs Induced by Cyclic Stretch in Control and HLHS iPSC-CMs

We next performed bulk RNA sequencing of stretched and unstretched tissues for all control and HLHS iPSC-CMs to identify transcriptional responses to stretch. For each line, unbiased principal component analysis (PCA) on the top 500 most variable genes revealed that cyclic stretch was a dominant regulator of gene expression, as captured by principal component 1 (PC1) in all control and HLHS lines (**Figure 4A**). Hierarchical clustering of the top 20 most significant differentially expressed genes (DEGs) separated tissues first by stretch condition (static vs stretched) and then by line (**Figure 4B**). Differential expression analysis across all six lines revealed 1176 significant genes (down-regulated: n = 839; up-regulated: n = 337; padj < 0.05, |log2FC| > 0.585), representing a core stretch-responsive transcriptional program that is preserved in both control and HLHS iPSC-CMs (**Figure 4C**). Genes that were notably downregulated due to stretch in all lines include fast twitch myosin isoform (*MYH6*), cholesterol efflux channel ATP-binding cassette-1 (*ABCA1*), Cytochrome P4501B1 (*CYP1B1*), and RNA Binding Motif (*RBM3*). Interestingly, 3 of the 4 HLHS patients showed high baseline expression of endocytosis gene *KIAA0319*, with HLHS 2 showing elevated expression after stretch. Notably upregulated genes in response to stretch include key ventricular maturation genes Homeobox Only Protein X (*HOPX*) and *MYL2*^27^, collagen matrix genes *COLQ* and *COL9A1*, chemokine *CXCL12*, and long non-coding RNA *OLMALINC*.

**Figure 4:**
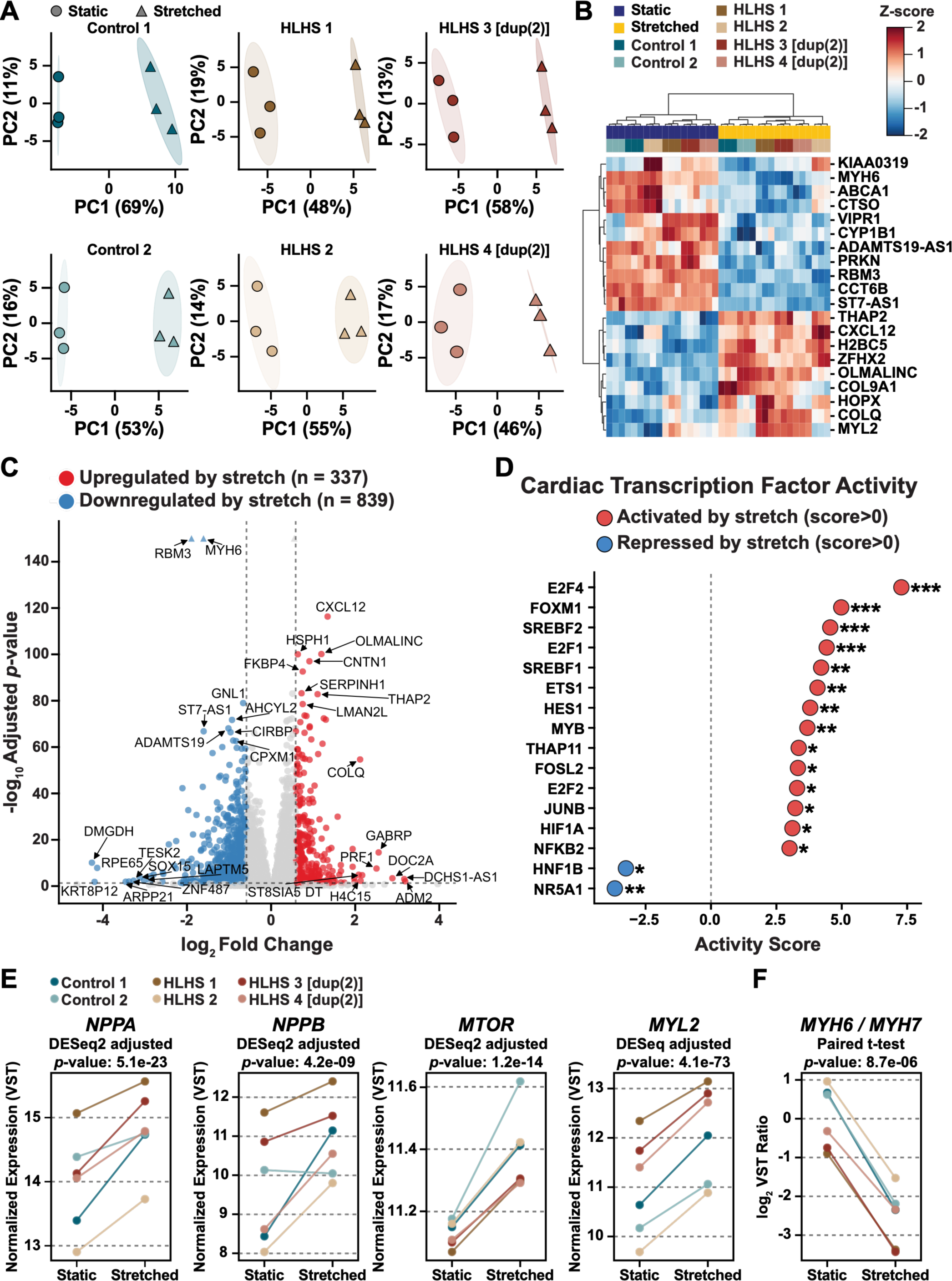
Stretch-induced remodeling of gene expression in control and HLHS iPSC-CMs. (A) Principal component analysis of the top 500 most variable genes for static (n=3, circles) and stretched (n=3, triangles) tissues for iPSC-CMs from each indicated line, analyzed independently. (B) Heatmap of the top 20 differentially expressed genes (*p-*adj < 0.05, |log2FC| ≥ 0.585). Each column is a tissue condition (see color legend) and each row is a gene, with z-score indicated by color bar. Euclidian hierarchical clustering shown in brackets. (C) Volcano plot of global differential gene expression between static and stretched tissues for all lines of iPSC-CMs,. Genes reaching significance thresholds (*p-adj* < 0.05, |log2FC| > 0.585) are indicated in red (upregulated, n=337) or blue (downregulated, n=839). Non-significant genes are indicated in grey. Axes are capped at |log2FC| = 6 and −log10(*p-adj*) = 150, with genes exceeding capped values indicated by triangular markers. (D) Transcription factor (TF) activity estimated from global stretch-induced gene expression for iPSC-CMs from all lines (stretched vs. static, log2 fold-change), restricted to cardiovascular protein-coding genes. Of 289 TFs, 16 reached significance (FDR<0.05). Red: increased activity due to stretch, blue: decreased activity due to stretch. Statistical significance determined from Benjamini–Hochberg FDR correction, \**p*<0.05, \*\**p*<0.01, \*\*\**p*<0.001. (E) Paired plots of expression of *NPPA*, *NPPB*, *MTOR,* and *MYL2.* The mean value of expression for static and stretched tissues for iPSC-CMs from each indicated line are plotted and connected to show trends. Adjusted *P*-values from DESeq2 (patient as a blocking factor, with Benjamini–Hochberg FDR correction) are indicated at the top of each plot. VST, variance-stabilizing transformation. (F) Paired plot of the log_2_ MYH6/MYH7 (α-/β-myosin heavy chain) expression ratio across static and stretched. *p*-value is calculated by 2-tail paired t-test on per-patient change in log_2_ MYH6/MYH7 from static to stretch.

To identify the regulatory programs underlying the transcriptional response to stretch, we inferred the transcription factor activity from the genome-wide stretch-induced expression signature, filtered using the cardiovascular disease atlas^28^. Sixteen transcription factors showed significant changes in activity due to stretch across all lines (*P < 0.05*; **Figure 4D**), including cell cycle regulators of the E2F family, lipid metabolic transcription factors (*SREBF1, SREBF2*), and hypoxia-responsive factors (*HIF1A*). Additionally, stretch induced activation of hallmark stretch-responsive genes, including natriuretic peptides A and B (*NPPA* and *NPPB*), mechanistic target of rapamycin (*mTOR*), and ventricular myosin light chain 2 (*MYL2)* (**Figure 4E**). The expected myosin isoform switching was also identified in all lines, marked by a decreased ratio of myosin heavy chain 6 to myosin heavy chain 7^29,30^ (**Figure 4F**). Together, these data indicate that many transcriptional responses to stretch are conserved in HLHS and control iPSC-CMs, suggesting that aspects of mechanosensing and mechanoadaptation are functioning normally in HLHS iPSC-CMs.

### Cyclic Stretch Induces Unique Transcriptional Programs in HLHS iPSC-CMs

Beyond the conserved stretch-responsive transcriptional program shared across all lines, analysis of the transcriptome on a per-line basis revealed that additional genes modified by stretch were largely patient-specific (Supplemental Figure 5A-C). This is generally expected given the variable genetic background of each patient and the sporadic nature of the disease. To delineate HLHS-specific differences in gene expression induced by stretch, we used DREAM analysis to identify stretch-induced genes that diverge between control and HLHS lines^31^ (Supplemental Figure 6A). In static tissues, 48 genes differed between control and HLHS lines, including known developmental/signaling regulators hairy and Enhancer of split-1 (*HES1*) and *NOTCH1*^32,33^, metabolic and lipid-handling gene Glycerol-3-phosphate acyltransferase, mitochondrial (*GPAM*), and cell division genes Centrosomal Protein of 57 kDa (*CEP57*) and Nucleoporin 37 (*NUP37*)^34–36^ (Supplemental Figure 6B) . In stretched tissues, 389 genes differed between control and HLHS lines. Of these, the top 30 divergence genes included sarcomeric and cytoskeletal genes, such as obscurin (*OBSCN*) and talin 2 (*TLN2*), and extracellular matrix genes, such as *COL2A1* and *COL9A1*^37,38^ (Supplemental Figure 6B). We next identified pathways enriched in the divergent stretch response and filtered them based on relevant keywords related to calcium, sarcomere, and inflammation. Several pathways related to calcium handling, contraction, and myofibrillogensis were significantly (FDR<0.05) enriched by stretch in control lines relative to HLHS lines (**Figure 5A**). By contrast, several pathways related to inflammatory signaling were significantly enriched by stretch in HLHS lines but not in control lines.

**Figure 5:**
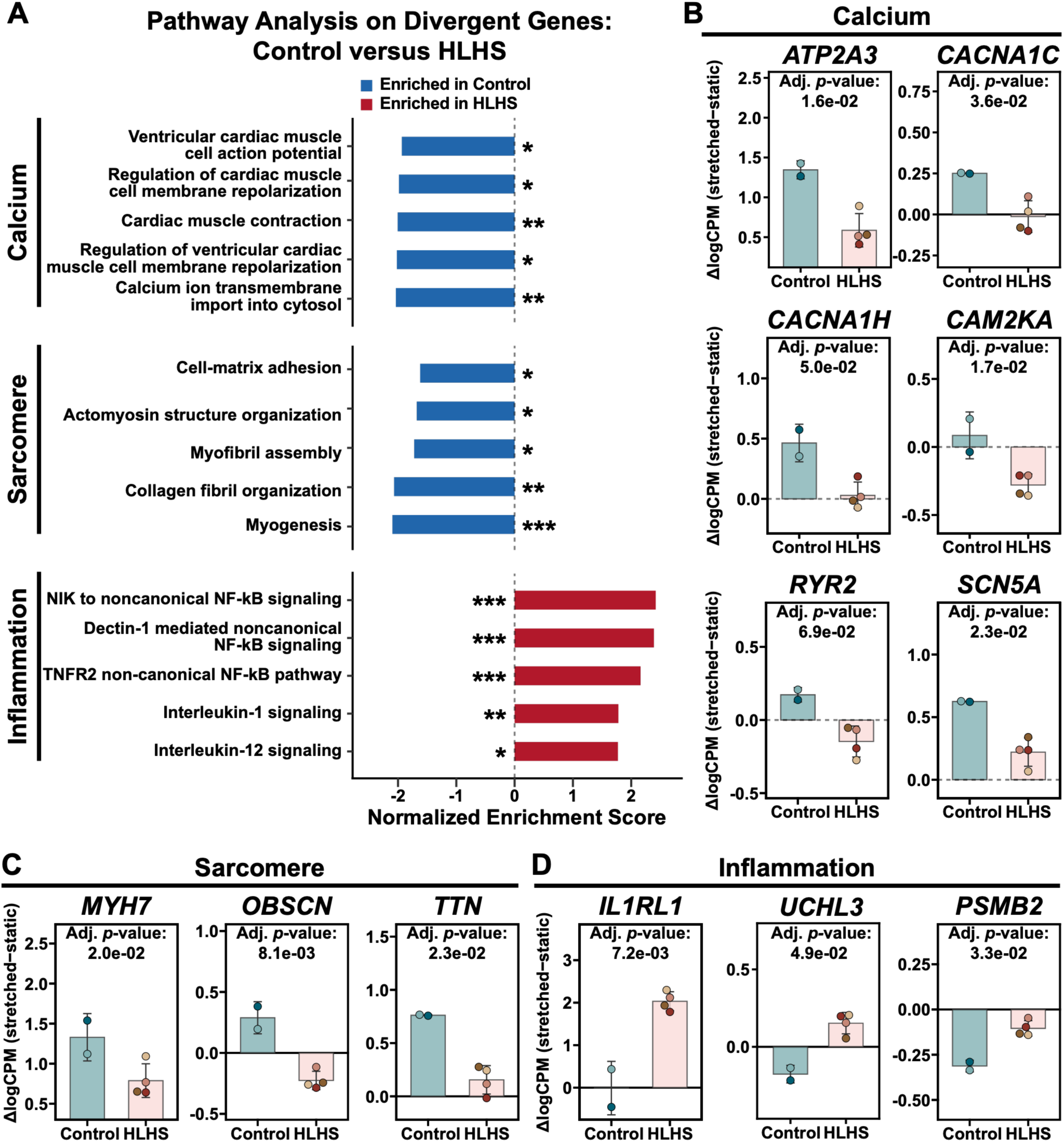
Divergent transcriptional responses to stretch in control and HLHS iPSC-CMs. (A) Gene set enrichment analysis (FDR<0.05) for pathways related to calcium, sarcomere, and inflammation. Bars indicate normalized enrichment scores. Blue indicates pathways enriched in control lines relative to HLHS lines due to stretch. Red indicates pathways enriched in HLHS lines relative to control lines due to stretch. *FDR<0.05, **FDR < 0.01, ***FDR < 0.001. Stretch-induced changes (ΔlogCPM, stretched – static) in select leading-edge genes related to (B) calcium, (C) sarcomere, and (D) inflammation. Bars indicate mean±SD, n=2 control lines (teal) and n=4 HLHS lines (pink). Adjusted *p*-values from the dream interaction coefficient are shown for each gene.

Next, we evaluated differences in specific hallmark genes related to pathways that were differentially enriched between control and HLHS lines. Related to calcium handling (**Figure 5B**), we found that *ATP2A3*, which encodes for sarcoplasmic reticulum Ca²⁺-ATPase 3 (SERCA3), and the voltage-gated calcium channels *CACNA1C* (L-type, Cav1.2) and *CACNA1H* (T-type, Cav3.2) were all significantly upregulated by stretch in control lines compared to HLHS lines. *SCN5A,* which encodes for the primary cardiac sodium channel Nav1.5, also increased significantly more in control lines compared to HLHS lines due to stretch. *CAMK2A*, which encodes for calmodulin-dependent protein kinase IIα, decreased due to stretch in HLHS lines but not control lines. *RYR2*, which encodes for ryanodine receptor 2, trended lower in HLHS lines but did not reach significance. Stretch-induced upregulation of sarcomere-related genes was also blunted in HLHS lines compared to control lines, with *MYH7* (β-myosin heavy chain), *OBSCN* (obscurin), and *TTN* (titin) all showing significantly higher expression due to stretch in control lines compared to HLHS lines (**Figure 5C**). Related to inflammation and stress responses, *IL1RL1* (ST2, the decoy receptor for IL-33) was not expressed in control lines but significantly increased in HLHS lines after stretch (FDR < 0.01) (**Figure 5D**). Lastly, *UCHL3* and *PSMB2* were significantly more suppressed by stretch in control lines compared to HLHS lines. These genes are part of the ubiquitin-proteosome system and their downregulation plays a role in activation of the NF-κB pathway^39^, a central inflammatory signaling system. Taken together, these pathway and gene-level analyses suggest that cyclic stretch induces physiological adaptation in calcium handling and contractility in control iPSC-CMs but not HLHS iPSC-CMs, which instead upregulate pathways related to stress and inflammation.

## Discussion

Our results demonstrate that HLHS iPSC-CMs in static culture exhibit mild impairments in sarcomere organization and calcium release compared to control iPSC-CMs. However, more striking differences were observed in the response to cyclic stretch. In both control lines, stretch consistently accelerated calcium release, a response observed in only one of four HLHS lines. This physiological divergence was mirrored transcriptionally, as control lines exhibited significantly greater stretch-induced up-regulation of several genes related to calcium release as well as sarcomere maturation compared to HLHS lines. Stretch also increased the expression of genes associated with inflammation and stress in HLHS lines compared to control lines. Together, these results indicate that HLHS iPSC-CMs have a maladaptive response to cyclic stretch, which may contribute to cardiac dysfunction in HLHS patients due to the essential role of mechanotransduction in normal cardiac development and physiology.

Similar to our results, several studies have demonstrated that cyclic stretch of iPSC-CMs induces maturation of the contractile cytoskeleton on the transcriptional level. This includes up-regulation of myosin regulatory light chain 2 (*MYL2*), a key component of sarcomeres in mature CMs^14,40^, and an increase in the ratio of *MYH7* to *MYH6*^41^, which encode for β-and α-myosin heavy chain, respectively. The switch from β- to α-myosin heavy chain is a hallmark of CM maturation and adaptation to mechanical load^40,42^. We observed similar trends in these genes across all control and HLHS lines, although several responses were blunted in HLHS compared to control, such as the stretch-induced increase in *MYH7*. *TTN*, which encodes titin, a key structural protein and mechanosensor in sarcomeres^43^ and established stretch-responsive gene^14^, was also strongly upregulated by stretch in control but not HLHS lines. Obscurin (*OBSCN),* which interacts with titin and anchors the sarcoplasmic reticulum with the sarcomere to enable efficient excitation-contraction coupling^43^, was up-regulated by stretch in control lines and down-regulated by stretch in HLHS lines. These examples demonstrate that HLHS iPSC-CMs have an impaired stretch-induced upregulation of several sarcomere-related genes. iPSC-CMs derived from patients with Down Syndrome also exhibit differential expression of sarcomere-related genes in response to cyclic stretch^44^, suggesting that defective cardiomyocyte mechanoadaptation may be a conserved mechanism in disorders associated with abnormal heart development.

Divergent stretch-induced transcriptional responses between control and HLHS iPSC-CMs were also observed in genes that govern intracellular calcium release. Cyclic stretch induced control but not HLHS lines to up-regulate expression of *CACNA1C. CACNA1C* encodes a subunit of the L-type calcium channel CaV1.2, which is primarily responsible for influx of calcium due to membrane depolarization and drives calcium induced-calcium release from the sarcoplasmic reticulum. A similar divergent expression pattern was observed with *CACNA1H*, which encodes for the low voltage-activated T-type calcium channel CaV3.2. CaV3.2 re-expression typically occurs selectively under pathological mechanical and neurohumoral stimulation via EGFR-dependent signaling^45^. Upregulation of *CACNA1H* would be expected to lower the depolarization threshold, facilitating earlier calcium induced-calcium release. *SCN5A,* which encodes for NaV1.5, the predominant voltage-gated sodium channel in CMs that facilitates the influx of sodium ions at the start of an action potential, upstream of calcium induced-calcium release, was also selectively upregulated by stretch in control but not HLHS lines. Thus, in response to stretch, HLHS iPSC-CMs failed to express several genes that encode ion channels that drive intracellular calcium release.

Consistent with the transcriptional data, our calcium transient data demonstrate that calcium influx in HLHS iPSC-CMs is dysfunctional, albeit with some line-to-line heterogeneity. In static culture, calcium influx was slower in all four HLHS lines compared to one of two control lines. In both control lines, stretch induced a significant acceleration of calcium influx, but this response was observed in only one of four HLHS lines. Calcium reuptake was unaffected by stretch in all lines, consistent with previous reports that stretch primarily affects intracellular calcium release but not reuptake^46^. Together, the functional and transcriptomic data suggest that HLHS iPSC-CMs have a distinctive impairment in adapting calcium influx to mechanical load.

Stretch also activated several transcriptional responses that were conserved between control and HLHS iPSC-CMs. All lines demonstrated enrichment of the E2F transcription factor family and *FOXM1*, which suggest universal engagement of hypertrophic growth and cell cycle re-entry programs in response to stretch^47–49^. *SREBF1/2*, which are key metabolic regulators that could drive fatty acid synthesis and lipogenesis in response to stretch^50,51^, and *ETS1*, *HIF1A,* and *NFKB2*, which could reflect a need for vascularization due to increased energetic demands^52–54^, were also upregulated by stretch in all control and HLHS lines. Other hallmark genes up-regulated by stretch, including the cardiac natriuretic peptides *NPPA* and *NPPB*^30^ and *MTOR*^55^, were also upregulated by stretch in all control and HLHS lines. These results indicate that HLHS iPSC-CMs do maintain some normal capacity to sense and adapt to mechanical loads, highlighting that dysfunctional remodeling of calcium influx is a specific physiological defect in these cells.

Our unbiased analysis of divergent pathways between control and HLHS iPSC-CMs also revealed that stretch activated stronger inflammatory responses in HLHS compared to control. Specifically, non-canonical NF-κB signaling pathways were enriched in HLHS lines after stretch. Over-activation of canonical NF-κB signaling is associated with heart failure and fibrosis, but less is known about the role of non-canonical NF-κB, though these pathways likely overlap in many downstream mechanisms^39,56^. Stretch also upregulated IL-1 and IL-12 signaling in HLHS lines, which have both been shown to play a role in downregulation of calcium gene expression and overload-induced cardiac inflammation, fibrosis, and ventricular dysfunction^57,58^. IL1RL1 was markedly upregulated by stretch in HLHS but not control lines. IL1RL1, also known as ST2, is a secreted decoy receptor for IL-33 and a well-established biomarker of cardiac stress and heart failure risk in congenital heart defect patients^59,60^. Together, these results suggest that cyclic stretch drives a coordinated inflammatory program in HLHS iPSC-CMs, indicative of a stress response that may contribute to contractile deficits, pathological hypertrophy, and progressive ventricular failure.

Collectively, this study demonstrates that iPSC-CMs from HLHS patients exhibit intrinsic defects in adapting calcium influx to mechanical load, a critical feature of normal cardiomyocyte physiology and mechanotransduction. Stretch also induces HLHS iPSC-CMs to activate inflammatory responses that could underlie many forms of ventricular dysfunction due to the broad effects of inflammation on myocardial tissue. Because patients with HLHS are exposed to shifting hemodynamic loads throughout the three staged surgical interventions^3^, an impaired ability of cardiomyocytes to adapt to mechanical loading could further weaken the overloaded single ventricle and exacerbate the risk of heart failure. Additionally, because cardiac morphogenesis is governed by both genetic programming and mechanical forces^61^, maladaptive responses to stretch could contribute to the abnormal development of the ventricle that is the defining feature of this disorder.

Our study also has important limitations. Given the inherent line-to-line genetic and transcriptional heterogeneity of iPSCs^62,63^, our results should be interpreted with caution due to the relatively low number of patient lines. Thus, expanding this study to more patient lines would enable more definitive conclusions and clearer genotype-phenotype correlations. We also used a standard ventricular CM differentiation protocol, which does not specifically distinguish left versus right CMs, an aspect that may be especially relevant to HLHS and should be considered in future studies. Other cardiac cell types, such as endocardial cells^6^, have also exhibited defects in HLHS but are not considered in this study due to our focus on CM-intrinsic defects.

## Acknowledgements

The authors would like to thank the lab of Justin Ichida for providing control patient lines and iPSC training and the Todd and Karen Wanek foundation at the Mayo Clinic for HLHS patient iPSC lines. We would also like to thank the Translational Imaging Center, the Molecular and Cell Biology Flow Cytometry core, and the John O’Brien Nanofabrication Laboratory at USC for providing facilities, equipment, and technical expertise. This work was funded by American Heart Association Transformative Project Award 25TPA1479668, NIH R01 HL153286, and Additional Ventures Expansion Award 1159088.

